# IW-Scoring: an Integrative Weighted Scoring framework for annotating and prioritizing genetic variations in the noncoding genome

**DOI:** 10.1101/163311

**Authors:** Jun Wang, Dayem Ullah, Claude Chelala

## Abstract

IW-Scoring represents a new Integrative Weighted Scoring model to annotate and prioritise functionally relevant noncoding variations. The pipeline integrates 11 popular algorithms and outperforms individual methods in three independent data sets, including variants in ClinVar database and GWAS studies, and cancer mutations. Using IW-Scoring, we located 11 recurrently mutated noncoding regions enriched for at least three functional mutations in 14 follicular lymphoma genomes, and validated 9 clusters (82%) in the International Cancer Genome Consortium cohort (n=36), including promoter and enhancer regions of *PAX5*. IW-Scoring offers greater versatility to identify trait and disease associated noncoding variants.

## Background

Over 98% of the human genome does not encode proteins. Despite its past reference as “junk DNA” when first discovered, noncoding sequences are now recognised as functionally important, possessing millions of regulatory elements and noncoding RNA genes. The annotation of potential regulatory sequences, through the Encyclopedia of DNA Elements (ENCODE) [1], Roadmap Epigenomics Consortium[2] and the FANTOM5 project [3, 4], is revolutionising our understanding of noncoding sequences and revealed that large stretches of the human genome (~80%) is evidently associated with DNA transcription to RNA, chromatin marks and other epigenomic elements. It has also established that many of the genetic variants associated with disease and diverse traits uncovered by genome-wide association studies (GWAS) are located within these noncoding regions, residing in or near ENCODE and Epigenome defined locations. However, compared to protein-coding regions, our understanding of noncoding regulatory elements remains poor.

Overall the significance of non-coding mutations remains an underexplored area of cancer genomics. With the rapid emergence of whole-genome and regulatory region targeted survey, many recurrent noncoding mutations have been identified in various cancer types. For example the prevalence of *TERT* promoter mutations has been established in melanoma [5, 6], gliomas and a subset of tumours in tissues with low rates of self-renewal [7]. Moreover, *TERT* promoter mutations significantly correlate with survival and disease recurrence in bladder cancer, demonstrating the clinical significance for noncoding mutations [8]. In chronic lymphocytic leukaemia (CLL), whole genome sequencing (WGS) of 150 tumour/normal pairs alongside with DNase seq and chromatin immunoprecipitation sequencing (ChIP-seq) identified recurrent mutations in the 3' UTR region of *NOTCH1* gene and in the active enhancer of *PAX5* [9]. In a subset of T cell acute lymphoblastic leukaemia (T-ALL), somatic mutations were found in the intergenic region to create MYB-binding motifs, which resulted in a super-enhancer upstream of the *TAL1* oncogene [10]. More recently, three significantly mutated promoters have been identified based on deep sequencing in 360 primary breast cancers, and more such regions remain to be discovered [11]. Whole-genome sequencing (WGS) data is increasingly available, especially through TCGA (The Cancer Genome Atlas) and ICGC (International Cancer Genome Consortium), prompting more pan-cancer style studies to look for significantly mutated regulatory elements across cancer types [12-15].

To systematically study these noncoding variants and assess their functional/pathogenic potential, they first require careful annotation, by determining the regulatory regions they map to, *e.g.*, ENCODE, Epigenome Roadmap and FANTOM5 defined elements, and their potential target genes. However, identifying potential noncoding pathogenic/driver/functional variants and distinguishing them from benign/passenger/non-functional variants, remains challenging, due to their abundance, cell/tissue type specificity and complex modes of action [16]. In the last few years, there have been many studies integrating all available noncoding annotation features to provide scores for the likely functional impact of noncoding variants. Many of these studies used machine-learning algorithms to develop classifiers integrating a range of annotations including regulatory features, conservation metrics, genic context and genome-wide properties to differentiate a set of disease-associated/deleterious variants from a set of benign/neutral variants in the model training, like CADD [17], GWAVA [18], FATHMM-MKL [19] and Genomiser [20]. Other methods, such as DeepSEA [21] and DeltaSVM [22], chose to directly learn regulatory sequence codes from large-scale chromatin-profiling data generated from ENCODE, while FitCons [23] opted to estimate the selective pressure for those functionally important genomic regions on the basis of patterns of polymorphism and divergence. Additional methods, like FunSeq2 [24] and Eigen [25], developed a weighted scoring system to combine the relative importance of various annotation features to separate functional and non-functional variants. The advantage of these weighted scoring methods is that they do not rely on any labelled gold-standard training data of disease-associated and putatively benign variants, which are often inaccurate and impractical for the extent of variants linked with GWAS and cancer studies.

Most of these methods above simply provide a continuous functional score for each variant without further information on the prediction accuracy and functional consequence in affected regulatory elements. A level of confidence or significance is also needed for each estimated score as these methods often score the same variants differently with varied performances across different data types. Thus, a class ensemble approach that combines the predictions of all these functional methods with a weighted scheme would offer a powerful approach to summarise multiple predictive evidences, increase specificity and rank variants, as this can generally improve the robustness and generalisability over a single scoring system [26].

To address these issues, here we explored the most commonly used functional annotation methods and scoring systems for genetic variations in the noncoding genome. Using an unsupervised spectral approach based on feature covariance, we propose a new Integrative Weighted Scoring system, IW-Scoring, amalgamating 11 existing functional scores into a single ‘ensemble-like’ algorithm that outperformed constituent methods in independent data sets. We then demonstrated its utility in identifying and prioritising functional variants in GWAS, expression quantitative trait locus (eQTL) and cancer studies.

## Results

### IW-Scoring framework

IW-Scoring consists of four top-level modules, including: 1) gene annotation, 2) regulatory annotation, 3) functional scoring, and 4) score integration and significance inference (**Fig. 1**). With queried variants as input, IW-Scoring framework uses the four modules to annotate and score variants, and rank noncoding ones based on their predicted functional significance. Within the gene annotation module, the queried variants are first annotated against Ensembl gene annotation to identify and filter out any variants that can potentially lead to non-synonymous changes for any transcript of a gene using our SNPnexus software [27]. For filtered noncoding variants, a gene centric view/summary is provided for within genes (e.g., synonymous, UTR and intronic), within 1 kb up- or downstream of genes, or intergenic. The next step is to further annotate noncoding variants against ENCODE/Epigenome/FANTOM5 defined regions along with Ensembl Regulatory Build annotation to identify overlapping regulatory elements of various chromatin (*e.g.*, DNase I), polymerase and histone (*e.g.*, H3K4me1/2/3 and H3K27ac) marks, transcription factor (TF) binding sites (*e.g.*, CTCF, FOXA1, NFKB, c-Myc, c-Jun, p300, etc) and predicted promoters/enhancers/TSS, with the supporting cell and tissue types also reported.

**Fig. 1.**
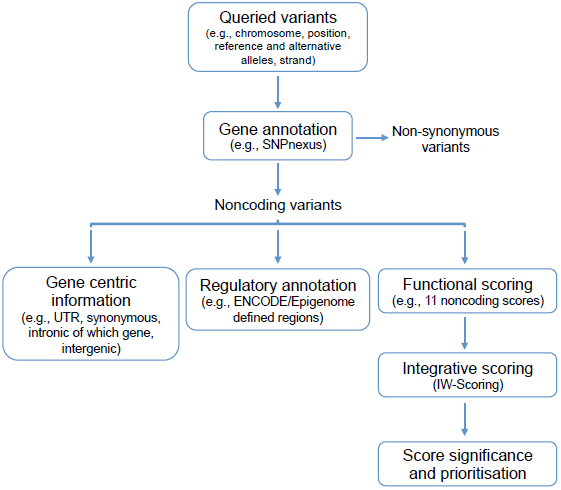
The IW-Scoring framework.

Next, noncoding functional scores of 11 different scoring systems derived from eight studies, including CADD v1.3 [17], DeepSEA [21], Eigen (Eigen and Eigen-PC) [25], fitCons [23], FunSeq2 [24], FATHMM-MKL [19], GWAVA (region, TSS and unmatched scores) [18] and ReMM [20] (**Additional file 1: Table S1**), are extracted for all noncoding variants allowing missing values for some scores. Finally, an integrative score, *i.e.*, IW-Scoring, is calculated for each variant as the weighted sum of all different available functional scores (see below for details), with associated statistical significance also derived.

### IW-Scoring for noncoding variants in training data set

We used an unsupervised spectral approach similar to that described by Eigen [25] to calculate associated weights for different scoring systems. Compared to the annotation resources used in Eigen/Eigen-PC, our IW-Scoring framework is designed specifically for noncoding variants, with the incorporation of a wider range of noncoding functional scores as described above. For the training set, we extracted all variants from the 1000 Genomes Project data set [28] that were not present in dbNSFP [29] v3.0, and were either 1) within 1 kb upstream of the gene start site and 1 kb downstream of the gene end site, or 2) within 5′ and 3′ UTR of a gene, or 3) synonymous variants in coding regions, leading to a total of 712,259 noncoding variants. Available functional scores were extracted and rescaled to calculate the covariance matrix (See **Methods** for details). Correlations among different functional scores for the training variants showed that scores from Eigen, DeepSEA, FATHMM noncoding, ReMM and CADD were more closely correlated, whereas GWAVA scores (*i.e.*, unmatched and TSS) were more similar to each other (**Fig. 2A**). Under the assumption of conditional independence among individual functional scores given the true state of a variant (either functional or non-functional), this correlation/covariance structure could be used to determine the weight for each scoring system when combined to differentiate variants. We are aware of the limitation of our assumption, as many functional scores tend to use similar information for prediction, thus likely to be correlated when scoring variants. However, our inclusion of a wide range of functional scores based on different subsets of regulatory annotations and different algorithms (either machine-learning classifier, or sequence pattern recognition, or sequence conservation or weighted summarising score) minimises the inter-dependency effect and ensure the functional scoring based on the most diverse evidences. It is worth noting that functional scores were available for >95% of training variants for all methods, except for FunSeq2 (84%), Eigen-PC (87%) and Eigen (91%), partially due to the missing values for chromosome X variants for Eigen (**Additional file 1: Table S1**).

**Fig. 2.**
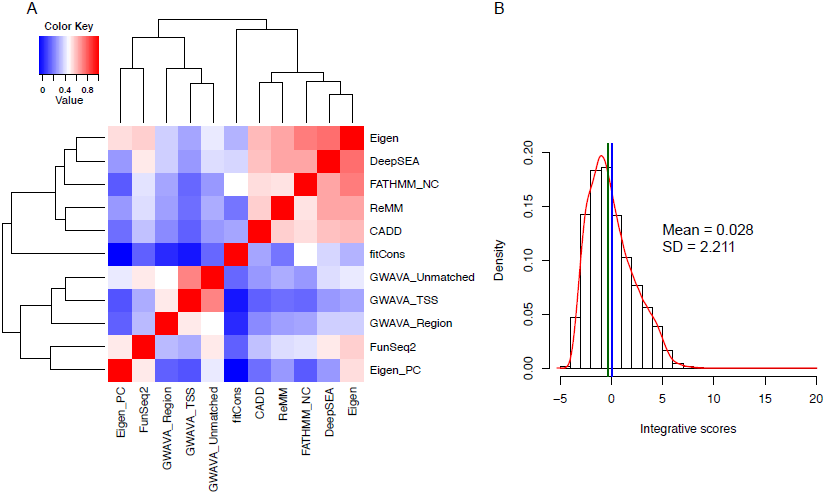
Correlations between 11 functional scores and the distribution of integrative scores of 712,259 noncoding variants in the training set. (A) Correlation matrix of 11 functional scores based on variants in the training set. (B) Distribution of integrative scores based on 11 scoring systems for all variants in the training set. The mean and standard deviation were shown. Mean and median values were indicated by the blue and green lines, respectively.

Using the eigendecomposition of the covariant matrix derived from the training set, we calculated the weights for different scores (**Additional file 1: Table S1**). Eigen, DeepSEA, FATHMM noncoding, ReMM and CADD had the highest weights, while fitCons, GWAVA TSS and Eigen-PC had the lowest weights, half of those for Eigen and DeepSEA. Our impression is that evolution and conservation based measures, like fitCons, may not be appropriate choice to score functional noncoding variants, as the regulatory elements are often lineage/species-specific [30]. We then computed the integrative scores for all training variants as the weighted sum of different functional scores (rescaled values) where the scores appeared to follow a lognormal distribution (mean of 0.028 and standard deviation of 2.211) with a long tail in the direction of positive values (**Fig. 2B**). This distribution could then be used to infer the statistical significance for all queried variants. We also compared the integrative scores amongst different feature types using the Ensembl Regulatory Build annotation, which revealed that variants in promoter regions had the highest scores, followed by those in promoter flanking regions and enhancers (**Additional file 2: Fig. S1**).

### Benchmark of IW-Scoring and comparison with other methods

Using three independent validation sets, we assessed the performance of IW-Scoring against other methods in differentiating pathogenic/functional noncoding variants from benign/non-functional ones.

### ClinVar pathogenic and benign noncoding variants

We selected noncoding single nucleotide variants (SNV) from the ClinVar database [31], including 3′/5′ UTR, intergenic upstream / downstream, intronic, and synonymous coding variants, resulting in in total 769 pathogenic SNVs (true positives) and 11,173 benign SNVs as control (true negatives). As shown in the average receiver operating characteristic (ROC) curves when comparing all 11 individual functional scores (**Fig. 3A**), ReMM, Eigen and FATHMM noncoding gave the best performances in differentiating pathogenic from benign variants with the area under the curve (AUC) values ranging 0.76 – 0.78 (**Additional file 3: Table S2**). The performances of DeepSEA and CADD were the next best with AUC of 0.72, whereas fitCons and GWAVA unmatched classifiers performed poorly (AUC = 0.58 and 0.62, respectively). We also performed the Wilcoxon rank-sum tests to compare functional scores between pathogenic and benign noncoding variants for each scoring system. Although the overall results were highly significant for all methods, *P* values from the tests were the most significant for ReMM (*P* = 1.5 × 10^−149^), FATHMM noncoding (*P*= 2.9 × 10^−149^) and Eigen (*P* = 2.5 × 10^−149^), while *P* values for fitCons (*P* = 6.0 × 10^−149^) and GWAVA unmatched (*P* = 7.3 × 10^−149^) were the least significant, consistent with the AUC results.

**Fig. 3.**
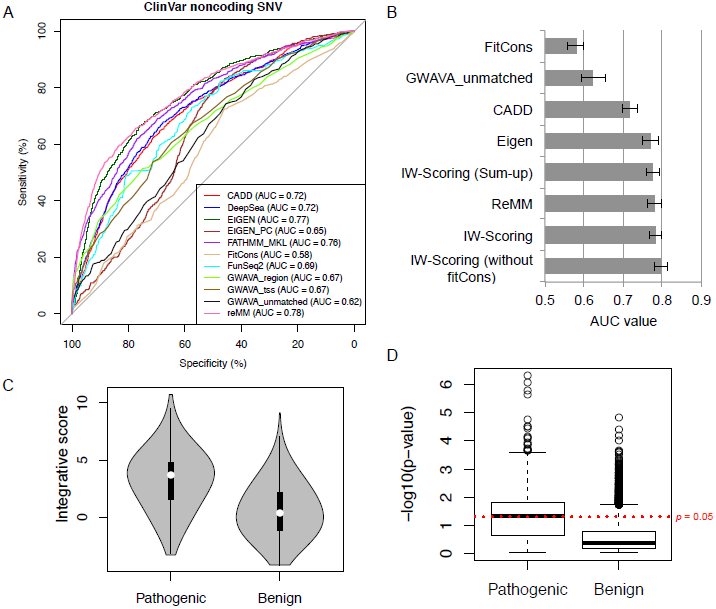
Comparison of the performances of different functional scores with IW-Scoring using the ClinVar pathogenic and benign noncoding variants. (A) ROC curves of the 11 individual functional scores with AUC values noted, measuring the accuracy to differentiate pathogenic and benign variants. (B) AUC values for IW-Scoring and selected individual scores to differentiate pathogenic and benign variants. 95% CI bars were also shown. (C) Violin plots of integrative scores (without fitCons) for ClinVar pathogenic and benign variants. (D) Boxplot of *p*-values associated with integrative scores (without fitCons) for pathogenic and benign variants.

We next calculated IW-Scoring values for all ClinVar pathogenic and benign noncoding variants. The AUC for the integrative scores was 0.78 (95% CI: 0.77 – 0.80), marginally higher than that of the best performing score above, ReMM (**Fig. 3B**). *P* value from the Wilcoxon rank-sum test was also the lowest for the IW-scores (*P* = 1.1 × 10^−149^). We also simply added up all 11 individual scores as the ‘sum-up’ scores, and the AUC for this was 0.78 (95% CI: 0.76 – 0.79), almost on a par with the weighted scores. As fitCons performed the worst out of all methods, we excluded fitCons from the training set and recalculated weights for all the remaining scores, followed by the calculation of IW-scores for all selected ClinVar noncoding variants. The AUC for this set of integrative scores to distinguish pathogenic from benign variants was 0.80 (95% CI: 0.78 – 0.81), the highest of all scores (**Fig. 3B**). This best performance was also supported by the most significant *P* value derived from the Wilcoxon rank-sum test (*P* = 1.5 × 10^−149^) (**Fig. 3C; Additional file 3: Table S2**). Furthermore, we also computed the significance *p*-values by comparing IW-scores with the lognormal distribution of those derived from the training set for selected ClinVar pathogenic and benign noncoding variants. *P*-values for pathogenic variants were much more significant overall than those for benign variants (combined *p*-value using Wilkinson’s method [15], *p* = 0.0003 for pathogenic variants and *p* = 0.15 for benign variants) (**Fig. 3D**). Within a queried variant set, IW-scores and associated *p*-values could be further used to rank and prioritise variants. It is worth noting that the performance of IW-Scoring did not improve when we excluded other methods with poor performances (*e.g.*, GWAVA unmatched) or just used the best performing methods only (*e.g.*, ReMM, Eigen and FATHMM noncoding). As shown in the distribution of scores across all tools for pathogenic and benign variants (**Additional file 4: Fig. S2**), IW-score (excluding fitCons) is close to taking the maximum of individual scores, increasing the specificity greatly. It is also important to note that IW-Scoring, along with ReMM and DeepSEA, resolved 100% of the selected ClinVar variants, with CADD and FATHMM noncoding scoring 98% variants, while three methods had scores for <90% selected variants, GWAVA (89%), Eigen-PC (86%) and FunSeq2 (only 55%) (**Additional file 3: Table S2**).

### GWAS significant noncoding SNPs

The second validation test data set we used was the noncoding trait-associated SNPs from the National Human Genome Research Institute (NHGRI)-EBI GWAS Catalog (http://www.ebi.ac.uk/gwas/home). We identified 19,797 GWAS significant noncoding SNPs, consisting of intergenic, intronic, synonymous and regulatory region variants. For negative ‘non-functional’ variants, we randomly selected more than twice the number of 1000 Genomes intergenic and intronic SNPs with minor allele frequency (MAF) > 5% in the population (n = 45,808). In general, the performances of all methods were much poorer for this data set when compared to ClinVar data set. This is probably because most of these GWAS significant SNPs are not truly causal, but in linkage disequilibrium (LD) with the genuine causal ones, which consisted of only 5% of all GWAS catalogued SNPs [32]. The AUCs for IW-Scoring and GWAVA unmatched classifiers were very comparable (0.591 and 0.595, respectively), producing the best results of all (**Fig. 4A**). Eigen-PC, DeepSEA, FunSeq2, GWAVA TSS and Eigen were the next best (AUC range 0.580 – 0.589), while ReMM and CADD performed relatively poorly (AUC, 0.533 and 0.536, respectively). FitCons appeared to be inadequate in handling this set of SNP data with AUC below 0.500 (**Fig. 4A; Additional file 5: Table S3**). Thus, we excluded fitCons from the integrative scores. In contrast with the ClinVar analysis, this exclusion did not improve the discriminating power of the integrative scores (AUC = 0.590) (**Fig. 4A)**. When comparing the *P* values from the Wilcoxon rank-sum test between GWAS significant and non-functional variants, the integrative scores led to the highest significance (*P* = 3.7 × 10^−149^) along with GWAVA unmatched, followed by Eigen-PC, DeepSEA, FunSeq2, GWAVA TSS and Eigen (**Additional file 5: Table S3**). Again, IW-Scoring was informative for all selected 65,605 variants, with ReMM, DeepSEA, FATHMM noncoding, FunSeq2 and CADD all scoring >95% variants. Eigen-PC was the only score system that had scores for <90% variants (89%) (**Additional file 5: Table S3)**.

**Fig. 4.**
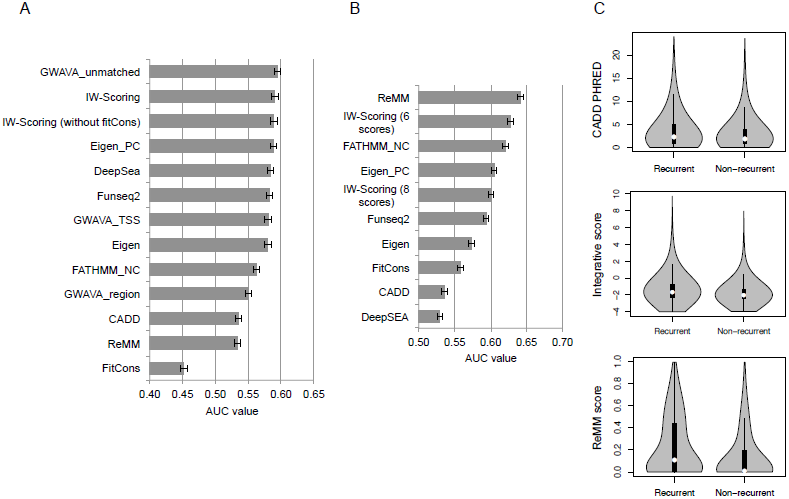
Comparison of the performances of different functional scores using the GWAS and COSMIC noncoding data sets. (A) AUC values for integrative scores and individual scores in the ability to differentiate between GWAS and randomly selected 1000G noncoding SNPs. 95% CI bars were also shown. (B) AUC values for integrative scores and individual scores in the ability to differentiate COSMIC recurrent from non-recurrent noncoding mutations. (C) Violin plots of functional scores between recurrent and non-recurrent noncoding mutations for CADD, integrative scores (six scores) and ReMM.

### Noncoding cancer mutations from COSMIC

Next, we compared the different scoring systems to determine their ability to differentiate potentially functional from non-functional noncoding cancer mutations in the COSMIC database [33]. As GWAVA only scores known SNPs, we omitted all three GWAVA scores from this test. Since the large-scale discovery of true functional noncoding mutations is still very much lacking, for potential “functional” ones, we identified all COSMIC noncoding mutations that had been identified in more than two samples and were also absent or present in less than 1% of samples in the 1000 Genome Project and NHLBI GO Exome Sequencing Project (ESP) data sets, restricting our analysis to 68,969 noncoding mutations. We further limited this set of mutations to those within the annotated regulatory regions only from ENCODE, Epigenome Roadmap and Ensembl Regulatory Build, leading to 34,813 potentially functional mutations as true positives. For the control set, we randomly selected ~4 million COSMIC non-recurrent noncoding mutations, and identified those with MAF larger than 1% in the 1000 Genome data set. We further excluded mutations within the annotated regulatory regions, leading to 57,866 noncoding mutations as the non-functional control. The AUC was the highest for ReMM (0.64), followed by FATHMM noncoding (0.62), Eigen-PC (0.60) and IW-Scoring of eight scoring systems (0.60) (**Fig. 4B; Additional file 6: Table S4**). CADD and DeepSEA appeared to perform the poorest in differentiating the selected recurrent from non-recurrent noncoding mutations, with AUC < 0.55. Thus, we excluded these two functional scores from IW-Scoring, and the new integrative scores based on the six remaining scoring systems significantly improved the discriminating power with an AUC of 0.63, which was only marginally lower than that for the best performing method, ReMM (**Fig. 4B; Additional file 6: Table S4**). In general, recurrent noncoding mutations within the regulatory regions had significantly higher functional scores on average than their non-recurrent counterpart for all scoring systems. The extent of this difference was the highest for ReMM and IW-Scoring compared to others (**Fig. 4C; Additional file 6: Table S4**). Again, Eigen and Eigen-PC resolved the lowest number of selected COSMIC variants.

In summary of the benchmark results across three data sets, the performance of IW-Scoring was stable and ranked consistently among the best performing methods. However, the performances of other methods appeared to vary greatly among data types.

### IW-Scoring usage and workflows

To further assist end-users to apply IW-Scoring to annotate and score their queried variants, here we discuss the standard usage of our framework. IW-Scoring contains separate workflows for scoring known and novel variants. For known variants, we provide two IW-scores with the associated *p*-values: one that aggregates scores from 11 functional scoring systems including fitCons, and the other excluding fitCons. This approach is useful for ranking and narrowing down variants indicated in GWAS and QTL studies, as well as rare known variants in hereditary diseases. As GWAVA only scores known variants, the workflow for novel variants excludes GWAVA scores from the aggregate calculation and provides two integrative scores: one that aggregates scores from the other eight scoring systems, and the other further excluding CADD and DeepSEA scores. This workflow is preferred for variants identified in cancer and other somatic diseases. In situations where users do not know which options to choose, they can use both workflows to generate IW-scores. The percentage of queried variants that are scored by GWAVA can further guide users in interpreting which IW-scores would be more reliable.

### Application of IW-Scoring to study noncoding variants from human data sets

Having assessed the performance of IW-Scoring and other available functional scores in a range of test data sets, next we aimed to use our scoring system to evaluate key noncoding variants derived from association mapping, eQTL and cancer studies.

### Disease SNPs near consensus motifs

We first assessed the functional significance for known disease-associated variants that fall near consensus motifs of transcription factor binding sites. We used candidate causal variants generated from Farh *et al.* [32], where transcription and *cis*-regulatory elements annotations for primary immune cells generated from Epigenome Roadmap Project were used and integrated. In this study, Farh *et al.* investigated the effects of disease SNPs in altering TF binding, and identified a notable AP-1 binding motif-disrupting SNP rs17293632 associated with Crohn’s disease. We extracted all nearby known variants, within 5 kb up- and downstream of rs17293632 (n=187), and calculated the IW-scores for them. Our results show that IW-score for the causal SNP rs17293632 is the highest (score = 9.48, *p* = 7.32 × 10^−13^) among all the SNPs investigated (**Fig. 5A**). rs17293632 is located within an intron of *SMAD3*, a region enriched for H3K4me1, H3K27ac, DNase I and TF ChIP-seq peaks, also with a higher degree of sequence conservation (**Fig. 5A**). The assessment of IW-scores further highlighted three additional SNPs with functional potential in this region, rs28514342 (5.47, *p* = 0.006), rs12324077 (4.98, *p* = 0.012) and rs193193326 (4.50, *p*= 0.020), which were all within 5 kb from the disease-associated rs17293632, thus likely to be in the same linkage disequilibrium (LD) block.

**Fig. 5.**
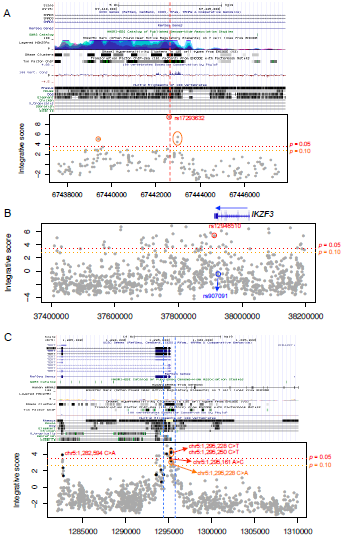
Application of IW-Scoring in scoring variants identified in GWAS, eQTL and cancer studies. (A) Disease SNPs near consensus motifs in *SMAD3* intronic region. Integrative scores (without fitCons) were shown for all nearby known SNPs along with the disease-associated candidates. (B) Disease SNPs with gene regulatory effects nearly 3' UTR of *IKZF3*. Integrative scores (without fitCons) were shown for 1,000 randomly selected known SNPs nearby, and two GWAS and eQTL candidates, circled with red and blue circles, respectively. (C) *TERT* promoter mutations and known SNPs in the region, with their integrative scores calculated. Black and grey dots denote somatic mutations and SNPs, respectively.

### Disease SNPs with gene regulatory effects

We also assessed the functional significance for disease SNPs with gene regulatory effects. Combining epigenome data and a data set mapping variants in peripheral blood gene expression, Farh *et al.* identified two eQTL SNPs in the *IKZF3* locus with independent effects on *IKZF3* expression. Based on the GWAS and eQTL significance, rs12946510 is associated with decreased *IKZF3* expression and increased multiple sclerosis (MS) risk, while rs907091 is associated with increased expression but with no association with MS risk. Interestingly, IW-score for the causal SNP rs12946510 is high (5.35, *p* = 0.007), whereas the IW-score for the eQTL SNP rs907091 with no disease effect is not significant (-0.105, *p* = 0.525), in line with the scores for all nearby non-functional SNPs (**Fig. 5B**). These results indicated that IW-scores are probably more sensitive to disease causative effects linked with phenotypic differences, rather than gene expression differences of nearby targeted genes. Therefore, by combining IW-scores and gene expression differences, we could further pinpoint those SNPs with both disease association and gene regulatory effects. *IKZF3* 3' downstream region is enriched for H3K27ac, DNase I peaks and TF binding sites; therefore, many variants in this region appeared to have higher IW-scores compared to those in other regions (**Fig. 5B**).

### Recurrent functional noncoding mutations in cancer

We assessed the functional significance of recurrent noncoding mutations identified in cancer, using *TERT* promoter mutations as the representative example. First, we identified somatic mutations curated in COSMIC and Huang *et al.* [34], as well as the known SNPs in this region, including ~15 kb up- and downstream flanking regions. In order to generate comparable scores between somatic mutations and known SNPs, we used the IW-Scoring novel variant workflow for both. Intriguingly, higher IW-scores were observed around the promoter region marked by strong DNase I signal and conserved TF binding sites, and gradually decreased to non-functional levels moving away from the promoter (**Fig. 5C**). For the two most frequently recurrent mutations, chr5:1,295,228 C>T and chr5:1,295,250 C>T (reverse strand, hg19), the integrative scores of eight scoring systems were 4.25 (*p* = 0.021) and 4.09 (*p* = 0.025), respectively. The scores for other less frequent mutations, chr5: 1,295,228 C>A and chr5:1,295,161 A>C (reverse strand), were 3.12 (*p* = 0.068) and 3.47 (*p* = 0.048). Encouragingly, this provided further confirmation of the accuracy and validity of our integrative approach for somatic variants. Interestingly, we also identified a silent mutation within a DNase I hypersensitive site, chr5:1,282,594 C>A (reverse strand), c.1719C>A, p.L573L, with a score of 3.88 (*p* = 0.032) with strong pathogenic potential (**Fig. 5C**).

### Landscape of noncoding mutations in follicular lymphoma

We next used IW-Scoring to investigate the landscape of noncoding mutations in cancer. We chose to examine WGS datasets in a haematological cancer, follicular lymphoma (FL) where the landscape of the coding mutations has been extensively studied recently [11, 35-37]. More than 90% of the tumours have mutations in genes encoding epigenetic regulators suggests that these tumours rely on epigenetic deregulation [38]. However, little is known about the repertoire of non-coding mutations in FL.

First we used a cohort of 14 WGS cancer samples from six FL patients from our previous study [11]. We calculated IW-scores for all 9,3078 unique noncoding somatic mutations. As expected, mutations in defined regulatory regions generally had significantly higher scores than those in non-regulatory regions (Wilcoxon test, *p*= 3.53 × 10^−49^, for Ensembl Regulatory Build; *p* = 2.71 × 10^−89^ for ENCODE annotation). For example, within Ensembl Regulatory Build defined regions, mutations in promoters had the highest IW-scores on average, followed by those in open chromatin and promoter flanking regions, while mutations in TF binding sites had relatively low scores (**Fig. 6A**). This is largely reflected by the occurrence of mutations in various ENCODE annotated regions (**Additional file 7: Fig. S3**). Among different ENCODE TF binding regions, mutations within *POU2F2, BCLAF1, FOXA2*, Ini1 and Brg1 binding sites seemed to have the highest pathogenic potential on average, further emphasising the importance of B-cell specific activator (*POU2F2*, also known as Oct-2) (mean IW-score, -0.47, compared to the mean of -1.15 for all mutations in TF binding regions), Bcl-2 associated protein (*BCLAF1*) (mean IW-score, -0.63), and chromatin modification and remodelling (*FOXA2*, Ini1/*SMARCB1*, and Brg1/*SMARCA4*, mean IW-score of -0.70, -0.83 and -0.85, respectively) in FL development (**Fig. 6B; Additional file 8: Table S5**).

**Fig. 6.**
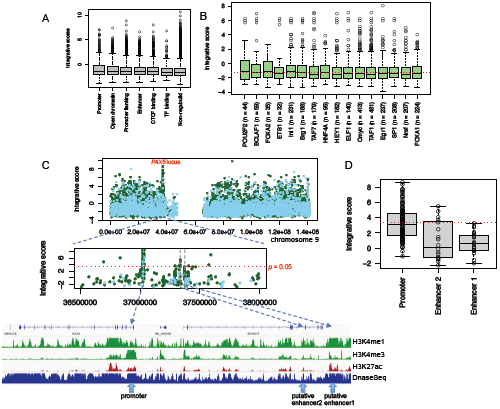
Integrative scores of somatic mutations in follicular lymphoma. (A) Integrative scores for mutations within Ensembl Regulatory Build annotated regions. (B) Integrative scores for mutations within top mutated TF binding sites. The mean value was shown by the red dotted line. (C) Recurrent mutations in *PAX5* locus. Integrative scores were shown for all noncoding mutations in whole chromosome 9 first. *PAX5* locus was further zoomed in with three mutations clusters further identified, *PAX5* promoter, putative enhance region 1 and 2. Mutations from ICGC FL samples were shown by solid green dots, while mutations from Okosun *et al.* were shown by solid light blue dots. Signals of epigenetic histone marks were for GM12878 cell line. (D) Integrative scores for mutations across three *PAX5* associated regions in boxplots. Red dotted line indicated the level of *p* = 0.05.

By using a *p* < 0.05 significance threshold for the IW-scores, this allowed the identification and refinement of the list of non-coding mutations to 1,375 (~1.5%) variants (**Additional file 9: Table S6)**. To highlight recurrent functional mutations in regulatory regions, we next examined the density of these candidate functional mutations and searched for regions where functional mutations were clustered within short inter-distance of each other (<10 kb). We identified 11 such clusters with enriched functional mutations (n ≥ 3) (**Additional file 9: Table S6**). Many of these are known targets of aberrant somatic hypermutations (aSHM) in B-cell lymphomas, such as *BCL2, BCL6, BCL7A, CXCR4* and *PAX5* [39, 40]. However, many recurrent functional mutations were also found outside the typical targeted regions of aSHM for these genes, *i.e.*, within ~2k bp downstream of transcription start sites (TSS). To what extent these mutations were associated with aSHM remains unclear.

We were able to validate these mutation clusters in an extended cohort of 36 FL patients derived from the ICGC Malignant Lymphoma Project where whole-genome simple somatic mutations (SSM) were available (ICGC Data Portal Release 23). Nine out of 11 clusters (82%) were also significantly enriched for functional mutations in this larger cohort (**Additional file 10: Table S7**), although many more mutational clusters enriched for functional mutations were discovered (data not shown). For example, we found in total 64 functional mutations in *PAX5* 5' upstream/UTR and first intron, and only 7 of these were within the usual aSHM target regions (**Additional file 11: Table S8**). The majority of these mutations were within *PAX5* first intron that contains an active promoter characterized by a DNase I hypersensitive site with strong H3K4me3 and H3K27ac signals in a lymphoblastoid B cell line GM12878 (**Fig. 6C**). Interestingly, we further identified two clusters of mutations ~300-330 kb upstream of the *PAX* gene. One cluster was within *ZCCHC7* intronic region with 5 potential functional mutations, and the other cluster of 24 mutations was located at the 3' downstream region of *ZCCHC7* with the most significant mutation as chr9: 3,7371,916 G>A (3.29, *p* = 0.058) (**Fig. 6C and 6D**). Both regions seemed to contain active enhancers marked by DNase I, H3K4me1 and H3K27ac peaks profiled in GM12878 (**Fig. 6C**). The latter cluster has been previously described in several B-cell related malignancies, indicating this region served as a *PAX5* enhancer [9]. By comparing FL mutations and their IW-scores among these three regions, our results further suggest that mutations in *PAX5* promoter were commonly more deleterious than those in enhancers (Wilcoxon test, *p*= 4.64 × 10^−149^, for promoter *vs.* enhancer region 2) (**Fig. 6C and 6D**). Future studies are required to further understand the gene expression regulatory effect and mechanism of these mutations in those regions.

## Discussion

The noncoding regions of the genome harbour a substantial fraction of total DNA sequence variations, and the functional contribution of these variants to complex traits, genetic diseases and tumourigenesis is still very poorly defined. Only a small fraction is believed to be truly functional and pathogenic. How to identify and prioritise these key functional driver events has become critical in the era of routine whole genome surveys and studies. We describe here an integrative approach, IW Scoring, for noncoding variant annotation and functional scoring. We showed that our approach outperforms most other functional scoring methods in differentiating functional/pathogenic from non-functional/benign variants in a variety of independent data sets: (1) validated clinically relevant variants, (2) GWAS significant variants, and (3)recurrent COSMIC cancer mutations. We also demonstrated its powerful application in identifying functional mutations in FL noncoding genome.

Our integrative approach has several advantages when compared to other available methods. First, it starts with embedded gene and regulatory annotation modules, allowing for easy access to the gene centric information and overlapping regulatory elements from a wide range of annotation resources, as well as available functional scores for queried variants. Second, it uses an unsupervised spectral approach to assign weights to available functional scores, and integrates these into a weighted sum. This approach yields the final scores and functional calls based on multiple evidences with a level of significance also derived to further improve the prioritisation. Thirdly, the nature of this integration meant that the performance of IW-scores was stable and ranked consistently among the best performing methods across all test variant data types. The performances of other methods, however, appeared to vary among data types, therefore, caution is needed to interpret these scores when different data types are studied and compared. In contrast with other tools, IW-Scoring benefitted by scoring the highest proportion of variants in all data sets via imputations based on interdependencies between variables. These advantages ensure that our method is versatile and can be applied to various studies from known SNPs, to rare germline variants and to somatic novel mutations.

A common feature of IW-Scoring and many other methods (e.g., CADD, Eigen and GWAVA) is that disease/phenotype and tissue/cell specificities with their related annotations were all combined into a single functional score. This technique significantly reduced the dimensionality of scoring output without having to produce scores for each tissue and phenotype, as well as for different chromatin/histone marks, making the data post-processing much easier and more straightforward especially for a large number of variants. However, IW-Scoring still allows for the functional variants associated with specific tissues, cells and features to be identified through the regulatory annotation module. This is currently lacking in many other methods, although some algorithms have chosen to focus on the identification of disease/tissue specific risk variants recently [22, 41]. Compared to most available methods, we believe our approach is optimally balanced between summarised and detailed evidences for the diverse range of users.

It could be argued that the approach we have adopted with IW-Scoring is similar to Eigen and Eigen-PC but it is worth pointing out that, our framework is specifically designed for noncoding variants, incorporating most recent and a wider range of noncoding annotation and functional features/scores. Via a vigorous weight learning process, strong weights were assigned to the block of closely correlated scores (Eigen, DeepSEA, FATHMM noncoding, ReMM and CADD), and the derived IW-Scoring significantly outperformed individual constituent scores (including Eigen and Eigen-PC) across various data sets, demonstrating the accuracy and validity of our approach. Such ensemble based approach with different estimated weights has been shown to perform better than any single component classifier [26], and has been widely used in various bioinformatics problems [41, 42]. The weighted integration technique based on the eigendecomposition of the covariance matrix also offers the flexibility to incorporate any other correlated genome-wide functional scores/features into the integrative scores. Many other integration approaches for continuous output can also be explored. For example, decision tree and random forest have been shown to perform well to predict the functional consequences of non-synonymous variants combining various scores [43]. Their performances on noncoding variants in comparison to our IW-Scoring approach are yet to be seen. However, as suggested by our results, incorporating more annotation scoring systems does not necessarily improve the performance of integrative scores, especially if these additional scoring systems are very much uncorrelated with the existing ones. Integrative scores can be mostly useful when combined with additional layers of genetic data, such as gene expression profiling by RNA-seq, to look for differential expression of target genes. This combinational approach has recently been applied to identify recurrent noncoding regulatory mutations in pancreatic cancer [44]. We are aware that IW-Scoring and all other algorithms are useful and powerful tools to prioritise mutations and associated genes from hundreds of thousands of noncoding mutations. However, these findings still need to be validated in the wet-lab to determine their true oncogenic potential.

In this age of WGS and diagnostics, we believe IW-Scoring offers great versatility in discovering noncoding disease causal variants. With the collection of a cohort of samples with the same disease or phenotype, one can identify recurrently mutated noncoding regions enriched for functional mutations predicted by IW-Scoring, which could drive the disease initiation and development, with or without the presence of coding drivers. This will certainly trigger the new wave of noncoding biomarker discovery.

## Conclusion

Here we describe IW-Scoring, an integrative weighted scoring framework for functional annotation and prioritisation of genetic noncoding variants. IW-Scoring shows consistent and superior performance in identifying potential functional/pathogenic variants and mutations when compared to individual constituent scores, across ClinVar, GWAS and COSMIC noncoding data sets. The four top-level modules in our approach offer dynamic outputs with a versatile range of gene and regulatory annotation features, as well as individual and integrative functional scores with associated significance. We demonstrate the utility of IW-Scoring in identifying functional noncoding variants and mutations in GWAS, eQTL and cancer studies. In particular, we harnessed the capacity of the algorithm to identify clusters enriched with functional or pathogenic mutations thereby providing a powerful tool to further decipher novel noncoding driver candidates in disease genomes. In the era of WGS and multi-omics studies, we propose that IW-Scoring can play an integral part in the search for significant regulatory variations in complex traits and diseases.

We have constructed a web interface for users to query into various noncoding annotation and scoring databases to annotate any list of known and novel variants, and to calculate IW-scores for them. This was made available at http://snp-nexus.org/IW-Scoring.

## Methods

### Data Sources

#### Gene coding annotation source

We use Ensembl (version 75) gene annotation model to annotate variants with SNPnexus and select noncoding variations. We also collect related gene mapping/effect information *e.g.*, synonymous, intronic, 5′ or 3′ UTR, up- or downstream (within 1k bp) of the gene. If variants are between genes, *i.e.*, intergenic, we collect their nearest up- and downstream genes and distances to them.

#### Noncoding annotation sources

We use ENCODE, Epigenome Roadmap and FANTOM5 for noncoding annotations. ENCODE and Epigenome Roadmap data sources were obtained from Ensembl BioMart (Version 75). Within ENCODE, annotated regions for DNase I, polymerase, various histone marks (*e.g.*, H3K4, H3K27, H3K36 and H3K9), TF binding across 19 cell lines are implemented for regulatory annotations. For Epigenome Roadmap, annotated regions for DNase I sites and up to 25 different histone marks for H1ESC and IMR90 cells are included. We also include Ensembl Regulatory Build annotation. For FANTOM5 regions, we collect overlapping CAGE peaks, predicted enhancers and TSS for each noncoding variant.

#### Noncoding variant scoring sources

We use 11 noncoding functional scores to build the IW-Scoring algorithm. These 11 scores were derived from eight studies, CADD, DeepSEA, Eigen, fitCons, FunSeq2, FATHMM-MKL, GWAVA and ReMM (**Additional file 1: Table S1**). For all these methods except DeepSEA, the pre-computed genome-wide score files with tabix indexes were downloaded. For DeepSEA, the original source code was obtained and run locally.

#### Construction of integrative noncoding scores

We used an unsupervised spectral approach similar to Eigen / Eigen-PC [25] to derive a weighted linear combination of individual scores. To achieve this, we first need to learn the weights for different functional scores incorporated.

#### Estimation of the weight for constituent scores

The weight estimation procedure is implemented as following:

1. *Training data set assembly*: a training data set of 712,259 noncoding variants was first assembled, consisting of variants within up-, downstream and UTR genic regions, as well as synonymous variants from the 1000 Genomes Project. We obtained functional scores from the 11 scoring systems, allowing missing values for certain scores. DeepSEA functional significance scores (*p*- values) were –log_2_ transformed to enable data integration.
2. *Data rescaling*: noncoding scores (continuous variables) were rescaled to have a mean of zero and a variance of one, individually. The minimum and maximum for the original and rescaled values of each scoring system were retained for the data transformation of queried data sets (see the section below).
3. *Covariance estimation*: a covariance matrix was calculated as pairwise correlations between any two of the 11 scoring systems. This allows variants with missing values for certain scores to be used when values from other methods are available. This ensures we estimate the weights based on all observed variants accurately.
4. *Weight estimation via eigendecomposition*: the estimated weights for the individual scoring systems were calculated using the eigendecomposition of the covariance matrix. The lead eigenvector (*i.e.*, the one with the greatest eigenvalue) was used and assigned to the corresponding scores as their weights.

We also calculated IW-scores, a weighted sum of rescaled values of different noncoding scores, for all training variants, in order to determine a distribution of integrative scores. We imputed missing values in the rescaled scoring matrix using the Amelia R package [45]. Amelia performs multiple imputation of missing data taking into account potential interdependencies between variables, combining the expectation-maximization with bootstrapping algorithm. The final imputed values were calculated as a mean of 10 imputations.

#### Calculating integrative scores and associated significance for queried variant sets

The IW-Scoring framework to calculate integrative scores for any set of queried variants is outlined below:

1. *Score extraction and missing value imputation:* similar to the training data processing, we first extract scores from different methods. For variants with missing values from certain scoring systems, imputations are implemented. Considering the usual small sample size for queried variant sets, we employ a strategy to merge them with a set of 100,000 variants randomly selected from the training set where all values are available, in order to increase the imputation accuracy. We then impute the missing values using Amelia based on the average of 10 imputations.
2. *Data rescaling*: we use the training data set as a reference for all queried variants to be rescaled to. For each scoring system, we rescale the values of queried variants to fit into the rescaled distribution of the training set based on the parameters (*e.g.*, the minimum and maximum) derived from the original and rescaled scores (at the training variant data recalling stage above). The rescaled values need to satisfy the following equation,

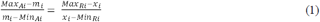

where Max_Ai_ and Min_Ai_, Max_Ri_ and Min_Ri_ are the maximum and minimum values of the original and rescaled scores for scoring system *i* in the training set, respectively. Variables *m*_*i*_ and *x*_*i*_ are the original and rescaled values of scoring system *i* for the queried variant. Thus, the solution to this equation of calculating *x*_*i*_ is

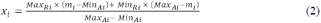
3. *Integrative scoring*: after rescaling, the integrative scores are calculated as the weighted sum of rescaled values of all scoring systems for queried variants, as

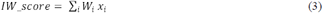

where *W*_*i*_ is the weight of scoring system *i* derived from the training set.
4. *Scoring significance and variant ranking*: the integrative scores for queried variants are compared to the lognormal distribution of all scores in the training set (*e.g.*, **Fig. 2B**) to determine the significance levels. Queried variants are further ranked based on IW-scores and *p*-values derived.

#### Performance comparison between integrative scores and other scores

We used noncoding variants from the ClinVar database version 2016/11/01 and National Human Genome Research Institute (NHGRI)-EBI GWAS Catalog version 2016/11/21. The performance of each scoring method was assessed using the average receiver operating characteristic (ROC) curves and the area under curve (AUC) using pROC R package [46]. For cancer data sets, we selected noncoding variants curated in the COSMIC database Version 79.

## Additional files

**Additional file 1: Table S1.** Summary of all 11 individual functional scores, with associated weights estimated (XLSX 53 kb)

**Additional file 2: Fig. S1.** Integrative scores for mutations within various Ensembl Regulatory Build annotation features for training variants (PDF 29 kb)

**Additional file 3: Table S2.** Assessment of the performances of various methods in the ClinVar data set (XLSX 41 kb)

**Additional file 4: Fig. S2.** Distribution of functional scores for selected ClinVar pathogenic and benign variants across all methods (PDF 281 kb)

**Additional file 5: Table S3.** Assessment of the performances of various methods in GWAS and 1000 Genome data sets (XLSX 49 kb)

**Additional file 6: Table S4.** Assessment of the performances of various methods in the COSMIC data set (XLSX 41 kb)

**Additional file 7: Fig. S3.** Integrative scores for mutations within various ENCODE annotated regions for FL noncoding mutations (PDF 29 kb)

**Additional file 8: Table S5.** FL mutations in TF binding sites (XLSX 41 kb)

**Additional file 9: Table S6.** 1,375 FL mutations with significant integrative scores of p < 0.05 (XLSX 147 kb)

**Additional file 10: Table S7.** Nine out of 11 clusters identified in Okosun et al. were validated in ICGC FL cohort (XLSX 12 kb)

**Additional file 11: Table S8.** Potential functional mutations in PAX5 promoter and first intron in Okosun et al. and ICGC FL cohort (XLSX 53 kb)

## Abbreviations

IW-Scoring: Integrative Weighted Scoring
ROC: average receiver operating characteristic curves
AUC: Area under the ROC curve
bp: base pair
ENCODE: The Encyclopedia of DNA Elements
FANTOM5: functional annotation of the mammalian genome
GWAS: genome-wide association study
TCGA: The Cancer Genome Atlas
ICGC: International Cancer Genome Consortium
WGS: whole genome sequencing
eQTL: expression quantitative trait locus
UTR: untranslated region
SNV: single nucleotide variant
SNP: single nucleotide polymorphism
COSMIC: catalogue of somatic mutations in cancer
FL: follicular lymphoma
aSHM: aberrant somatic hypermutations

## Acknowledgements

We thank Jude Fitzgibbon, Jessica Okosun and Trevor Graham for their valuable comments.

## Availability of data and materials

IW-Scoring has been made available via http://snp-nexus.org/IW-Scoring.

## Authors’ contributions

JW conceived the study. JW, DU and CC designed and performed the experiments. JW wrote the manuscript. JW, DU and CC edited and reviewed the manuscript. All authors read and approved the final manuscript.

## Consent for publication

Not applicable.

## Ethics approval and consent to participate

Not applicable.

